# A modular gate system for autonomous control of rodent behavior

**DOI:** 10.1101/2024.11.22.624912

**Authors:** Adam W. Lester, Gurnoor Kaur, Nadira Djafri, Manu S. Madhav

## Abstract

Rodent mazes have been used for decades to study the neural basis of behavior. Advancements in rapid prototyping techniques and access to affordable electronics allows laboratories with sufficient expertise in engineering and programming to customize and construct maze apparatuses and behavioral tasks, thereby increasing the ability of their studies to answer specific scientific questions. We designed and built a rodent gate system that lowers this bar of expertise even further. The NC4gate system is a robust mechanical design that can be built using low-cost hardware and execute thousands of cycles before maintenance. Up to 512 gates can be controlled using a single computer. Users can control the gates interactively using a Python-based graphical interface and programmatically using an extensible API. We hope that the open-source hardware / software and extensive documentation enables laboratories to build these affordable and robust gates and seamlessly incorporate automatic behavior control into their existing or new rodent tasks.

**Significance Statement:** Rodent mazes are used by thousands of laboratories and research institutions across the world to study learning and memory, as well as the effects of pharmacological, genetic and environmental manipulations. Ideally, maze and task designs should be customized to the scientific questions at hand. It is challenging, however, for many laboratories to build, program, and operate custom mazes, requiring them instead to rely on expensive and proprietary commercial solutions. The most complex components of most mazes are moving gates that restrict and direct rodent behavior. Here we provide the open-source hardware and software for a gate system that is extensible, affordable and robust, removing this critical barrier to customized mazes.

## Introduction

Rodent mazes are widely used tools in behavioral neuroscience research to study learning, memory, and spatial navigation in rodents, particularly mice and rats. Researchers use these mazes to assess cognitive abilities, explore the effects of genetic manipulations or drug treatments, and investigate neural mechanisms underlying navigation and memory formation in rodent models. By utilizing rodent mazes, scientists gain valuable insights into the intricate workings of the brain and behavior in mammals, further contributing to our understanding of various neurological disorders and cognitive processes.

Rodent mazes can vary in complexity and design, including T-mazes, Y-mazes, radial arm mazes, and Morris water mazes (Bimonte-Nelson, 2015). These mazes typically consist of chambers or alleys in which cues, rewards or punishments are delivered. Movement of rats is restricted using either manually operated doors, gates or blocks. Each trial of an experiment is typically split into phases, each of which is defined by a particular behavioral constraint or stimulus. Traditionally, a researcher manually observes the rodent behavior, transitions between phases and delivers stimuli / rewards. The presence of the experimenter can influence rodent behavior, creating an uncontrolled and often unmeasured variable that can bias performance and slow down the pace of training. Remote and even autonomous operation and monitoring of maze systems is now possible due to advances in hardware, software and computing, which significantly mitigates confounding factors related to experimenter interactions.

Standard mazes are available to buy from commercial vendors, although the hardware and software are often proprietary and expensive. These mazes can often only run a limited set of predefined tasks and are not amenable to hardware or software modifications due to proprietary components. The availability of low-cost electronics, rapid prototyping methods such as laser cutting and 3D printing, as well as traditional machining services, allows labs to manufacture their own hardware tailored to their needs. Labs that choose this route still face significant challenges - building robust and well-executed maze hardware requires expertise in animal behavior, iterative design principles, as well as hardware / software development and integration. While there are many open science tools for experimental tracking (Dunn et al., 2021; Karashchuk et al., 2021; Mathis et al., 2018; Pereira et al., 2022), control (Akam et al., 2022; Lopes et al., 2015; Quigley et al., 2009), and analyses (Buccino et al., 2020; Rübel et al., 2022; Viejo et al., 2023), there are comparatively fewer open hardware solutions (Aguiar et al., 2020; Birtalan et al., 2020; Cano-Ferrer et al., 2023; Krynitsky et al., 2020; Nguyen et al., 2016; Oh et al., 2017; Ozgur et al., 2023) that can be easily integrated into existing / proposed experimental apparatuses.

We have designed the NC4gate, a modular remote operated gate system that can be integrated into maze apparatuses. The gate is low-cost, uses standard hardware and affordable manufacturing techniques, and is extremely robust in operation. The low-profile design allows multiple gates to be packed side-by-side. Up to 512 gates can be controlled using a single PC interface and microcontroller. The system can be remotely controlled manually using a GUI or autonomously using an API. In this manuscript, we provide the hardware design, PCB schematics, firmware and software to enable rodent behavioral researchers to adopt and adapt this technology into their experimental systems.

## Materials and Methods

The NC4gate system has three components – mechanical, electrical and software. Design files and assembly instructions are available on our Open Science Framework page (https://osf.io/uy7ez). These designs are made available under the CERN Open Hardware License – Weakly Reciprocal (CERN-OHL-W).

### Mechanical design

The NC4gate is built from flat components that are designed to be mated together primarily using snap-fits and a minimum number of mechanical fasteners. These flat components are fabricated from laser-cut POM-C acetal, a thermoplastic that provides high stiffness, low friction and excellent dimensional stability. Sheets of acetal (3.175 mm / 0.125 in thick; Robertson Plastics, Surrey, BC, Canada) were precisely cut using a laser cutter, a technology often available in academic machine shops or through affordable third-party services. Acetal can also be cut using machine tools such as Computer Numeric Control (CNC) mills, however, we recommend using a laser cutter due to the intricacy of the cuts required, which cannot be achieved with even very small diameter end mills. While other plastics such as acrylic are dimensionally stable and can also be laser-cut, we do not recommend them due to their brittleness, which may cause cracks or breaks after repeated operations. 10 acetal parts are laser cut for each gate. They can be arranged to fit in a single (11.875” x 15.875” or 300 × 400 mm) sheet that can fit the bed of most small laser cutters, including the one we use (Rayjet 30W; Trotec, Mississauga, ON, Canada). This allows components for each gate to be cut in a single run, saving manufacturing time and material costs.

A DC gear motor (37GB520, 12 V, 50 rpm, 37 × 60 mm, 100:1 metal gear, 50 kg.cm torque, 6 mm shaft dia.; available as FIT0492-A, DFRobot, Shanghai, China) drives the actuation of the gate’s wall panel (Fig. 1). The use of a DC gear motor, as compared to stepper motors, provides a lower cost solution and reduces auditory and EM noise interference. The motor is connected to the wall panel (100 mm wide) through a motor linkage and a pin- and-slot mechanism (Fig. 1). As the motor rotates, the linkage moves a follower inside the gate slot, driving the gate up and down. We manufactured this linkage in aluminum using water-jet machining. 3D-printed or laser cut versions of this arm were tested – however, they were not strong enough to endure repeated operation in our limited tests. There may be other materials or technologies that can allow equally robust operation using more accessible fabrication services. The specific mechanism used here provides a gate height of 180 mm, which is close to the height of the complete assembly. Linear actuators, in contrast, would need to be at least double the gate height (360 mm) for comparable operation. Limit switches mounted on either end of the gate slot register the limits of travel (Fig. 1).

**Figure 1:**
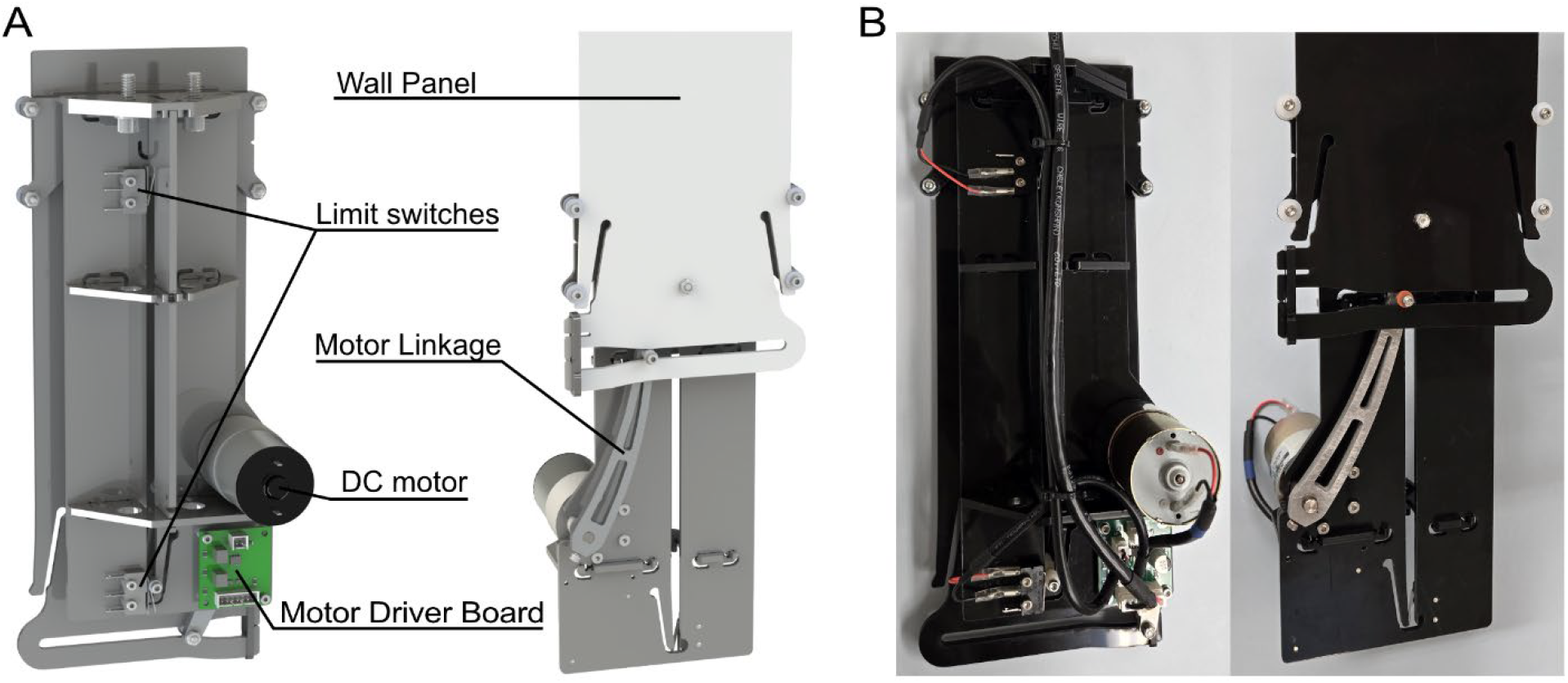
NC4gate module. (A) Rendering showing the back and front of the gate module and the major components. The Wall Panel moves up and down with a total travel of 180 mm. The limit switches detect the upper and lower limits of wall travel. The Motor Linkage actuates wall movement by translating from radial to linear motion using a pin-in-slot mechanism and is connected to a 12 V 50 RPM geared DC motor. The custom NC4 Motor Driver Board drives the motor via PWM through the H-bridge and relays the switch signals to the NC4 Cypress Board. (B) Pictures showing an actual assembled NC4gate module complete with the required wiring.

In addition to the 10 laser cut parts, motor, linkage, and limit switches, we use standard hardware for the various fasteners, spacers and rollers of the assembly. A full bill of materials is available along with detailed assembly instructions on our design page (Lester et al., 2024). Computer-Aided Design (CAD) models of the entire gate assembly were designed in Solidworks (v2022, Dassault Systèmes, Vélizy-Villacoublay, France) and are also provided. A 3D printed jig is recommended to drill holes to attach the linkage to the motor shaft, and a model of this is also available on the design page (Lester et al., 2024).

The gate module measures approximately 117 × 86 × 290 mm when lowered. This low-profile design allows for gate modules to be closely spaced for configurations requiring numerous gates.

With multiple gates in operation, failure of components can cause experimental delays and training issues. We tested N=5 gate modules with cyclic up / down runs with an interval of 5 seconds; they lasted thousands of cycles before failure (see Results). These figures ensure that the mechanical components of each gate will last through months of intensive operation. When components do fail, their low-cost and accessibility means that they will be quick to manufacture and replace so that experiments can quickly resume.

The gate module can be mounted to a horizontal surface using two standard fasteners and a slot approximately 102 mm long and 5 mm wide in the surface provides enough clearance for the wall panel of the gate to pass through. A cutting template for the slot and mounting holes is available on the design page (Lester et al., 2024).

### Electrical design

Several electrical components for the operation of these gates are custom-designed. Printed-circuit board (PCB) schematics were developed using Altium Designer (v23.11.1, Altium Limited, La Jolla, CA, U.S.A.) and are available on our design page (Lester et al., 2024). PCBs were manufactured by PCBWay (Hanzhou, Zhejiang, China) – however, other PCB manufacturers also have similar capabilities. Interconnecting cables use non-proprietary connectors. Cable specifications are provided on the design page (Lester et al., 2024). Other electronics components were sourced from online vendors such as DigiKey and Mouser and links are provided for the specific components and vendors we used in bill of materials (Lester et al., 2024).

### NC4 Motor Driver Board

Each NC4gate includes a single NC4 Motor Driver Board (Fig. 1; Fig 2.). The primary motor driver chip (DRV8870DDAR; Texas Instruments, Dallas, TX, U.S.A.) amplifies two 5V pulse-width modulation (PWM) signals to drive the DC gear motor in the up and down directions. The motor is connected to the NC4 Motor Driver Board through a wire with a 2-pin receptacle (0353120260; Molex, Lisle, IL, USA). The NC4 Motor Driver Board is also connected to two limit switches (MS0850501F025C1A; E-Switch, Minneapolis, MN, USA) mounted on either end of the slot in the NC4gate, through 2x 2-pin receptacles (Fig. 2; B02B-PASK; JST, Waukegan, IL, USA). Two transistors (SMMBT3906LT1G; Onsemi, Phoenix, AZ, USA) on the NC4 Motor Driver Board cut off power to the motor when either limit switch is triggered, ensuring safe operation and rapid stopping when the walls travel limit is reached. The NC4 Motor Driver Board communicates to an external system, such as the NC4 Cypress Board (see below) or stand-alone microcontroller, through a 7-pin receptacle (0353120760; Molex, Lisle, IL, USA), which provides power: motor voltage (12V, >0.68 A), logic voltage (5V, 100 mA) and ground, inputs: PWM down and PWM down, and outputs: upper and lower limit switch states. Any 5V microcontroller with at least at least 8192 bytes of RAM that can output two PWM signals and monitor two digital inputs (e.g. Arduino Uno, Arduino SA, Lugano, Switzerland) can be used to control a gate module. Depending on number of PWM channels and digital Input/Output (I/O) pins, multiple gates can be controlled from a single microcontroller. For instance, an Arduino Uno R3 can control 3 gates, and an Arduino Mega 2560 Rev3 can control 7 gates. Integrating the NC4 Cypress Board into the system, however, vastly expands this capability.

**Figure 2:**
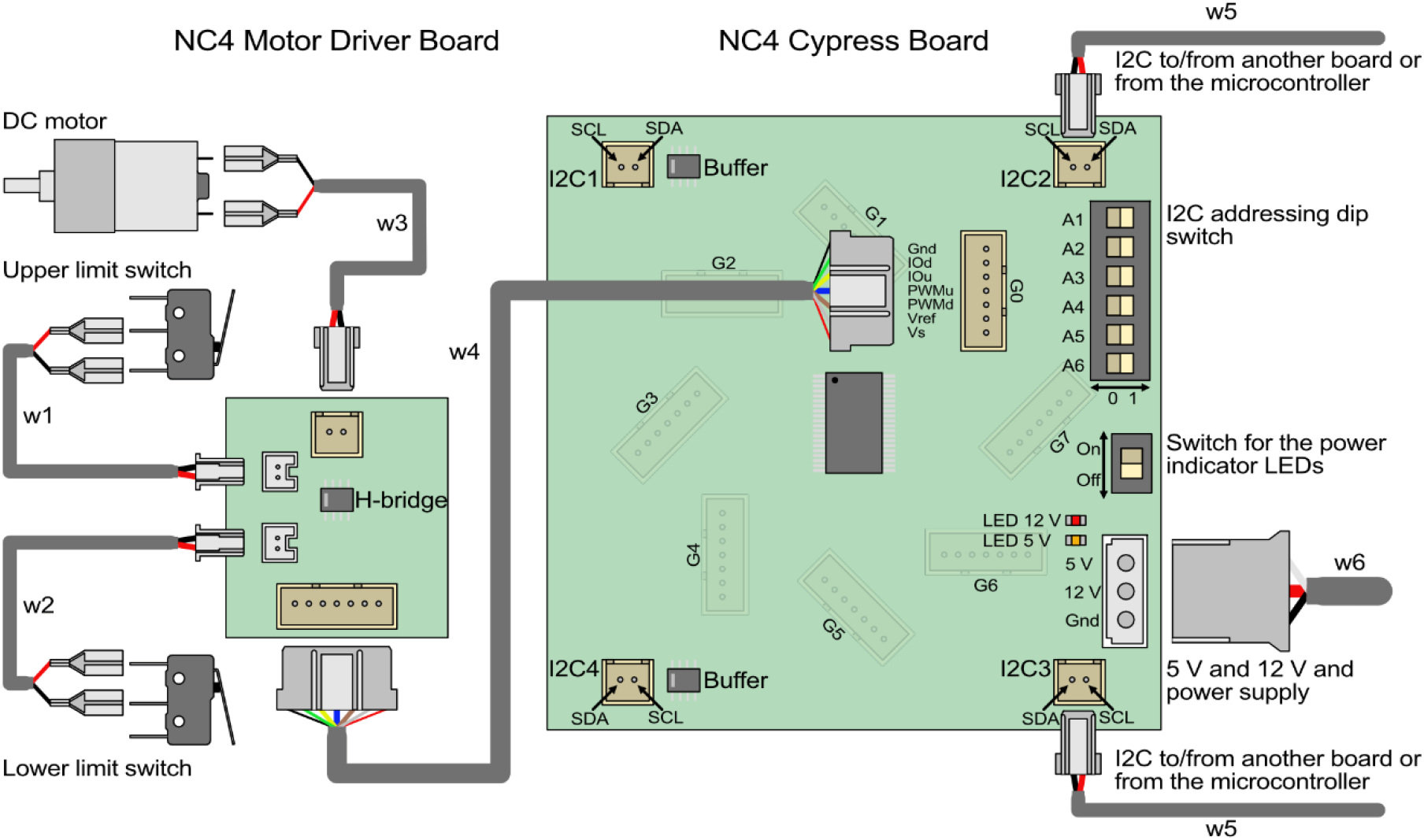
Electronics and wiring diagram. Illustration showing the major components of the custom NC4 Motor Driver Board and NC4 Cypress Board. Each NC4 gate includes a NC4 Motor Driver Board connected to a single 12 V geared DC motor as well as 2 limit switches, which register the gates position. The NC4 Motor Driver Board can be connected to one of eight 7-pin connectors on the NC4 Cypress Board. The NC4 Cypress Board requires 12 V and 5 V power and multiple can be daisy chained together via the I2C connectors and controlled through a single microcontroller.

### NC4 Cypress Board

The NC4 Cypress Board is built around the CY8C9540A (Infineon Technologies, Neubiberg, Germany), a chip that allows the configuration of up to 40 pins as inputs, outputs, or PWMs, and communicates to a microcontroller through the Inter-Integrated Circuit (I2C) protocol. The 7-bit address of each NC4 Cypress Board can be easily changed using 6 binary switches integrated into the board, with the first bit being fixed at 0. Thus, up to 2^6^ = 64 boards can be addressed on the I2C bus. We configured this chip to control the up/down PWM and monitor the up/down limit switch states of 8 gates (32 channels). Thus, a single microcontroller can theoretically control up to 64 × 8 = 512 gates; we have tested simultaneous operation of up to 60 gates using 9 NC4 Cypress Boards. An LED indicates powered status of the board but can be turned off using a switch for operation in low-light conditions.

I2C communication at 100 KHz (Standard mode) is limited to a few meters due to the buildup of bus capacitance, which is limited to 400 pF as per the I2C specification (NXP Semiconductors, 2021). To counter this, the NC4 Cypress Board features a pair of buffer chips (P82B715; Texas Instruments, Dallas, TX, U.S.A.) that isolate the capacitance of each chip, and increase the bus capacitance limit between Cypress chips to 3000 pF. Four 2-pin I2C connectors, SDA/SCL (35312-0260; Molex, Lisle, IL, USA) are provided on each NC4 Cypress Board. Two of these connectors can be used at a time. Using horizontal or vertical connections puts the NC4 Cypress Board into the common I2C bus, thus sharing its bus capacitance, while connecting diagonally isolates and branches the board into a new I2C bus. In this way, multiple buffered circuits can be created allowing groups of gates and their associated NC4 Cypress Boards to be positioned over longer distances. We tested the capacitive tolerance of the circuit to verify these properties (see Results).

The NC4 Cypress Board has a 3-pin power receptacle for 12 V, 5V and ground (350210-1; TE Connectivity, Schaffhausen, Switzerland). The NC4 Cypress Board connects to the NC4 Motor Driver Board of each gate through eight 7-pin receptacles (0353120760; Molex, Lisle, IL, USA).

### Microcontroller, power and PC interface

As mentioned above, most 5V-logic microcontrollers can be used to control a few gate modules. Once extended with the NC4 Cypress Board, we recommend using the Arduino Mega 2560 R3 (Arduino S.R.L., Monza, MB, Italy), which has enough SRAM (8 KB) for storing the firmware that we provide for gate operation and monitoring. This microcontroller connects to a PC through a standard USB serial interface.

A 12 V power supply is required for motor operation – each motor is rated for 0.68 A, and we recommend scaling the capacity of the power supply according to either the total number of gates, or maximum number of gates that will operate simultaneously. 5V power can be drawn from the microcontroller – we found this to be sufficient to operate 9 NC4 Cypress boards supporting 60 gates simultaneously. However, an independent 5V power supply is recommended for larger numbers of boards / gates. A bus bar can be used to deliver 12V and 5V power to each NC4 Cypress Board from a single source.

### Wiring

Depending on if the system necessitates using the NC4 Cypress Board, or users instead choose to use just a microcontroller, two wiring topologies are possible: Microcontroller <–> NC4 Motor Driver Board; Microcontroller <–> NC4 Cypress Board <–> NC4 Motor Driver Board.

Operation of the gate module requires two 2-pin cables (lengths of 36 cm for UP switch and 17 cm for DOWN switch, w1 and w2 in Fig. 2) between each limit switch and the NC4 Motor Driver Board, which terminate on the board by a header (PAP-02V-S; JST, Waukegan, IL, USA). A 2-pin cable (12 cm length, w3 in Fig. 2) connects to the motor with spade connector and terminates on the NC4 Motor Driver Board by a header (0351550200; Molex, Lisle, IL, USA). A 7-pin cable (40 cm, w4 in Fig. 2) terminated on the NC4 Motor Driver Board by a header (0351550700; Molex, Lisle, IL, USA) and the other end of this cable can use the same connector, if using an NC4 Cypress Board, or can be connected directly to the relevant header pins of a microcontroller.

Operation of the NC4 Cypress Board additionally requires 2-pin I2C cables (w5 in Fig. 2), terminated on the board by a header (0351550200; Molex, Lisle, IL, USA) and connected to either another board or a microcontroller. Additionally, a 3-pin power cable (14 AWG recommended, w6 in Fig. 2) is required, terminated by a header (1-480303-0; TE Connectivity, Schaffhausen, Switzerland).

We use cable assemblies pre-crimped with the appropriate headers from Cloom (Shijiazhuang Cloom Tech Ltd., Shijiazhuang, Hebei, China), although these wires can also be easily crimped and assembled in-house. Holes are provided on the gate module panels that can be used to cleanly route and secure wires using zip ties so that they don’t interfere with gate operation (Fig. 1).

### Software design

#### Arduino firmware

We have written firmware for the Arduino 2560 R3 that allows it to interface with multiple NC4 Cypress Boards, monitor the status of the gates, and control their operation. The code is available in a public github repository (https://github.com/NC4Lab/NC4gate), with instructions for flashing and testing a controller. The firmware communicates to the PC software using a USB serial interface. Template code is also provided in the repository to connect a gate directly to an Arduino for smaller operations.

#### PC software

Our public github repository hosts a cross-platform python-based API and Qt-based GUI that interacts with the Arduino firmware and allows a user to manually operate multiple gates connected to multiple NC4 Cypress Boards. The GUI automatically polls available NC4 Cypress Boards connected to the Arduino I2C bus and populates interfaces for gates connected to each of these boards. While the GUI allows for set up, testing and manual operation of gates, the API can be customized to run fully autonomous experiments. We aim to integrate the API with experimental control packages such as Bonsai (Lopes et al., 2015) to provide an easier way for animal behavior laboratories to integrate our gate into their standard operation.

### Code accessibility

The Arduino firmware and PC-based GUI for operation of the NC4Gate system are publicly available in our github repository: https://github.com/NC4Lab/NC4gate

## Results

Repairing and replacing components can significantly affect experimental timelines and the validity of the scientific results. While our gate module is easily repairable and replaceable, we specifically went through numerous design and testing iterations to ensure reliability and robustness in operation. We built 5 copies of the latest gate design (v1.1, Fig. 3) and tested each in the following manner.

**Figure 3:**
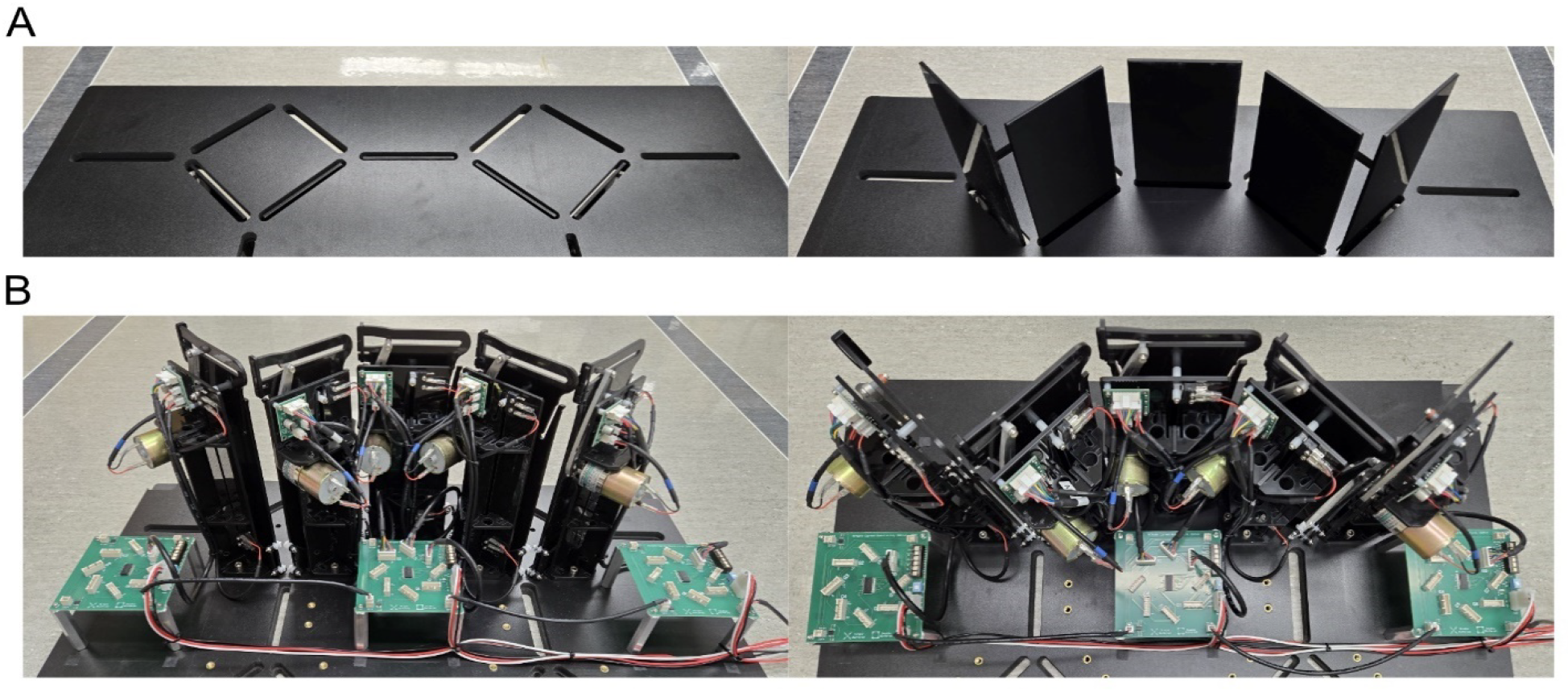
Example configuration utilizing 3 NC4 Cypress Board to and 5 NC4gates. (A) Pictures showing the five gates in their lowered and raised configuration. (B) Pictures showing the underside of the platform with all components and wiring.

### Timing

We timed the actuation of 5 NC4gate modules by connecting them to an NC4 Cypress Board using the test setup shown in Figure 3. Over 50 cycles, we measured the actuation duration, i.e. time between lower limit switch release and upper limit switch press when the gate is moving upwards, and upper limit switch release and lower limit switch press when the gate is moving downwards (Fig. 4). On average, upward movements took 578 +/-17 ms, and downward movements took 531 +/-13 ms. These durations are in practice perfectly acceptable for control of rodent behavior. Individual walls also had consistent durations, characterized by their low standard deviation over cycles (1, 5, 5, 1, 2 ms during upward movement, and 3, 5, 1, 2, 2 ms during downward movement).

**Figure 4:**
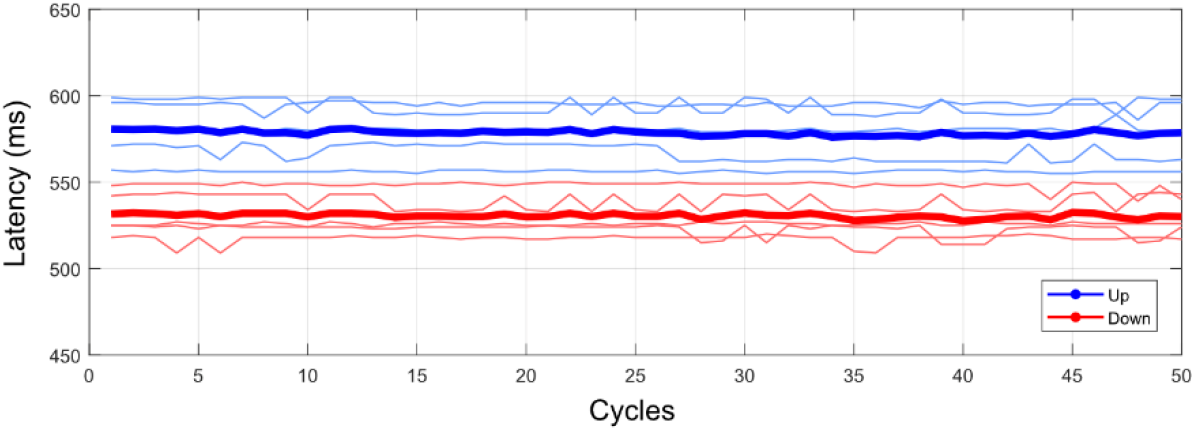
NC4gate operation timing. The latency between upward wall movements (blue) and downward movements (red) are plotted for 5 wall modules over 50 cycles. Latencies are plotted for each wall (thin lines) as well as averages (thick lines).

### Stress Test

After performing the timing tests, the DC motors, Motor Linkages, and limit switches were replaced on the 5 NC4gates. To stress test the gate’s mechanical robustness, gates were tested without the NC4 Cypress Board. Instead, an Arduino Mega was connected directly to the NC4 Motor Driver Board IO lines and a custom testing script was used to cycle the boards up and down with a cycle time of 5 seconds. The software counted the number of cycles until the eventual failure of a gate to trigger a limit switch. The results of these tests are summarized in Table 1. Over the course of continued testing, the gates failed at cycles 5258, 17227, 100000, 36307, 41271. Note that one of the gates reached 100000 cycles without failing, at which point the testing was ended. The observed critical failure modes included: mechanical seizure of the geared DC motor for 2 gates; the Retainer Clip unseating from the Wall Panel, which caused the separation of the Motor Linkage from the Wall Panel for 1 gate; and faulty wiring of the test setup, which was unrelated to the gate’s functionality for one gate. Additionally, 3 gates displayed loosening of the fastener between the Motor Linkage and the DC motor shaft.

**Table 1:** Stress Test Results for the NC4gates. This table summarizes the results of stress tests performed for 5 NC4gates. Mean and standard deviations (SD) are provided. Each gate was tested in parallel to determine the maximum number of operational cycles (Total Cycles) before failure occurred. The Duration of each test is provided in hours, minutes and seconds (HH:MM:SS) to highlight the total runtime. Failures encountered during the tests are categorized as either Non-Critical Failures, which represent minor issues not significantly impacting system performance, or Critical Failures, which led to complete failure of the gates to actuate. The non-critical failures primarily involved loose linkage connections, while critical failures included mechanical seizures of the DC motors and, in one case, wiring issues in the test setup unrelated to the NC4gates functionality.

| Wall Number | Total Cycles | Duration (minutes) | Duration (HH:MM:SS) | Non-Critical Failures | Critical Failures |
| --- | --- | --- | --- | --- | --- |
| 0 | 5258 | 438 | 07:18:10 | None | Mechanical seizure of DC motor |
| 1 | 17227 | 1436 | 23:55:35 | Motor Linkage connection loosened | Test wiring failure unrelated to component functionality |
| 2 | 100000 | 8333 | 138:53:20 | None | None |
| 3 | 36307 | 3026 | 50:25:35 | Motor Linkage connection loosened | Separation of the Motor Linkage from the Wall Panel due to Retainer Clip unseating |
| 4 | 41271 | 3439 | 57:19:15 | Motor Linkage connection loosened | Mechanical seizure of DC motor |
| Mean | 40013 | 3334 | 55:34:23 |  |  |
| SD | 32683 | 2723 | 45:23:35 |  |  |

### Capacitance

The I2C bus at standard speed (100 KHz) can only tolerate a capacitance of 400 pF as per the standard (NXP Semiconductors, 2021). Capacitance between the SDA and SCL lines increases proportional to the length of the cable. Therefore, there is an inherent limitation (∼50cm using a conservative estimate of 8 pF/cm of cable) on the spatial extent of the gate modules relative to each other, which is clearly not ideal for a many behavioral apparatuses. To overcome this limitation, we added a pair of P82B715 buffer chips to each board, with the appropriate pullup resistors on the input and output sides. Any two of the four I2C connectors on each Cypress board can be used as its input and output. When diagonally opposite connectors are used (Fig. 5A), only the input buffer on the board is engaged. While this will isolate the board itself from the I2C line, capacitances on the input and output side will not be isolated from each other (Fig. 5B) and will be additive. When vertically or horizontally opposite input and output connectors are used (Fig. 5C), both input and output buffers on the board are engaged, isolating the input and output capacitances from each other (Fig. 5D). This double buffering, however, can create a minor latency in I2C signalling (∼250 ns) which could become noticeable cumulatively. Since both options are available on each board, experimenters can opt to use single buffering on closely-spaced boards or double buffering between further-spaced boards to achieve the best compromise between signal integrity and latency.

**Figure 5:**
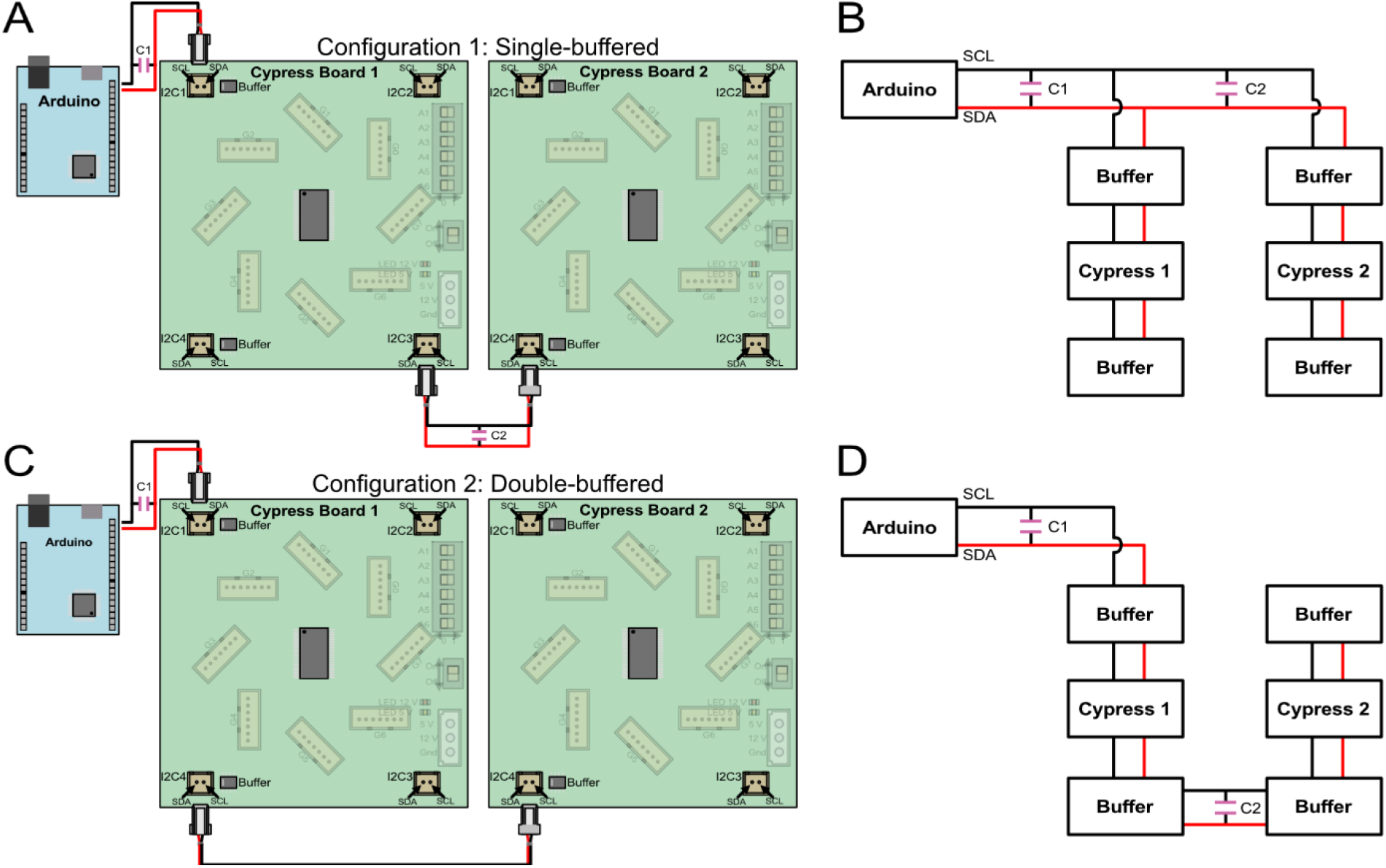
Capacitance testing setup. One Arduino was used to address two Cypress boards. (A) In the single-buffered configuration, diagonally opposite I2C connectors were used as input and output for Cypress board 1. Additional capacitors C1 and C2 were used to add line capacitance on the input and output sides (B) Schematic for A. Only one buffer on the input side is engaged, resulting in C1 and C2 being parallel (additive) on a common I2C line. (C) In the double-buffered configuration, vertically opposite connectors were used as input and output for Cypress board 1, though horizontally opposite connectors can also be used. (D) Schematic for C. In this configuration, both buffers on Cypress board 1 are engaged, resulting in C1 and C2 being disconnected on independently buffered circuits.

Using an LCR meter, we tested the capacitance of the Cypress board, between the SDA and SCL lines, which was ∼52.5 pF, consistent over frequency ranges from 10 KHz to 10 MHz. We also tested the capacitive tolerance of the Cypress boards in both the single- and double-buffered configurations. Figure 5A shows the diagram and Figure 5B shown the schematic of the single-buffered test. We increased input (C1) and output (C2) capacitances to board 1 in 100 pF increments, while testing whether both boards could be addressed. Since C1 and C2 are in parallel (Figure 5B), increasing either will produce the same result. Both boards were able to be addressed when C1+C2 <= 2000 pF.

Figure 5C shows the diagram and Figure 5C shown the schematic of the double-buffered test. Now C1 and C2 are independent, and we similarly increased each of them in 100 pF increments in separate tests. When C1 increased beyond 2000 pF, I2C communication failed to both boards. When C2 was increased beyond 3100 pF, I2C communication was retained to board 1, while board 2 lost communication.

We can conclude that we have a capacitive tolerance of ∼2000 pF between the Arduino and the first double-buffered Cypress board (2.5 m length at 8 pF/cm) and ∼3000 pF between each pair of double-buffered Cypress boards (3.75 m at 8 pF/cm). These figures are well within the spatial extents of typical mazes, allowing for flexible deployment of the NC4gate system.

## Discussion

The NC4gate is a low-cost (∼$50 USD) apparatus that enables robust and repeatable experimental control in a variety of rodent mazes. Behavioral science laboratories often rely on proprietary hardware / software packages from manufacturers that are expensive, sometimes justifiably so due to high R&D costs and low volumes. Standard mazes such as the T-maze and Y-maze are available commercially – however, building a custom maze still requires significant technical and system integration know-how. Additionally, such mazes are specialized for a single route topology. Paired with open-source behavioral tracking, reward delivery and experiment control systems, laboratories will be able to use our gate and controller boards to quickly prototype custom training paradigms. Fueled by availability and affordability of programmable electronics and open-source software, social behavior and homecage-based training and monitoring paradigms have become popular in recent years (Grieco et al., 2021; Jhuang et al., 2012; Murphy et al., 2016; Singh et al., 2019). Open hardware, i.e. electromechanical designs that are replicable, easily manufactured and reliable, integrate these electronics and software architectures into usable packages that are accessible by laboratories without advanced technical expertise, that nevertheless would benefit from custom experimental hardware.

The NC4gate modules allow placement and operation of only a few gates through direct microcontroller interface, and up to 512 gates through daisy-chained control boards. This allows the system to be used in a variety of scenarios, from manual remote control of a few gates by a human operator, to training of rodent behaviour in a fully-automated manner. We are currently operating a maze apparatus composed of 60 gates.

The modular and low-cost nature of the hardware enables labs to try building and operating an NC4gate before fully deploying it in the experimental systems. The open hardware / software and design files (Lester et al., 2024) allows laboratories with only limited access to fabrication resources to build their own NC4gates from readily available stock materials and hardware. Additionally, for more advanced users with some degree of familiarity with Solidworks modeling, the provided assembly files can be modified to customize certain parameters of the module (Lester et al., 2024), e.g. to reduce travel, change gate dimensions, or adjust speed of operation, etc. Use of laser cut parts and standard hardware for manufacturing enables these designs to be iteratively tested and improved.

In a typical rodent maze task, each trial typically involves 1-3 gate actuation cycles. Even our least durable gate lasted 5258 cycles, representing several months of operation. Failures were caused by the DC motor, retaining clip, wiring, and loosening fasteners, all issues easy to prevent by quick inspection or to fix with the replacement of inexpensive parts. Notably, the DC motor was the source of failure for the gate which achieved the fewest cycles of 5258. Consistent with our observations from other tests performed, the DC motors tend to be the primary point of failure in the system. To date, we have been operating a maze system with 60 NC4gate modules almost daily for 10 months with only a few retaining clips needing to be replaced. We have also observed that rats can be habituated to the moving gates in one or two sessions, are not injured on the rare occasion that a gate accidentally actuates beneath them.

From the manuscript and the design page (Lester et al., 2024), the reader may have noticed that metric and imperial sizes are intermixed in materials and fasteners. This is due to our aim of using the lowest cost and most accessible components in North America. However, substitutions (e.g. 0.125 in to 3 mm material, 4-40 to M3 screws) might be made in some cases with minimal design changes, without affecting functionality. For best results, however, we encourage users to use the same hardware specified in our bill of materials, which includes quantities, part numbers, and vendors for every component used (Lester et al., 2024).

Remote-operated or automated maze systems enabled by the NC4gate can significantly reduce experimenter intervention and associated confounds in rodent behaviour. The flat surface of the raised gate panels offers the ability to display projected or printed visual cues on them. The narrow profile of the gate modules allows them to be placed close together, reducing the overall size requirements of a given maze system.

Cost, lower EM noise and negligible idle power draw are three of the major reasons why we chose to use DC motors as compared to stepper motors in the gates. We have tested the gate modules with our extracellular electrophysiological recording system, which includes a SpikeGadgets Main Control Unit and 128-channel Horizontal Headstage (SpikeGadgets, Inc., San Diego, CA, USA) and Cambridge NeuroTech 64-channel silicon probes (Cambridge NeuroTech, Cambridge, UK). Using our system, we found that the DC gear motors operating beneath the rat do not introduce any detectable EM noise that could interfere with neural data collection. Introducing electromagnetic shielding under the maze surface (between the rodent and motor / electronics) and proper grounding can help alleviate EM noise in experimental apparatuses where this becomes an issue.

The NC4gate modules, in conjunction with the NC4 Cypress Board, utilize I2C communication to simplify the hardware requirements significantly, reducing wiring and enabling control of up to 512 gates with a single microcontroller for large-scale applications. We have incorporated I2C buffer chips to the NC4 Cypress Board design to improve I2C signal fidelity over the larger spatial distances required for most experimental apparatus. Still, the communication bandwidth limitation of Standard-Mode I2C (100 Kbits/s) (NXP Semiconductors, 2021) may introduce delays in control. These effects are minimal with our largest gate count (60) but may become significant with larger numbers of gates. This limitation is imposed by the CY8C9540A chip, which cannot operate in Fast mode (400 kbits/s), Fast mode+ (1 Mbit/s) or High-speed modes (1.7 Mbit/s). Port expander chips that operate in these modes could be used to enable Fast mode capabilities if required.

The NC4gate system offers a low-cost, scalable solution for automated control of rodent maze apparatuses that can address the demands of diverse experimental setups. By leveraging carefully designed, documented and tested open-source hardware and software, the NC4gate minimizes the need for technical expertise from the end user, while still allowing for varying degrees of customization by those with more advanced fabrication or programming skills. By designing the system with an eye towards high durability across thousands of cycles and ease of maintenance, robust operation with minimal downtime is achievable for both for small-scale and large applications. Use of an automated control approach increases behavioral throughput and reproducibility of environmental manipulations while reducing experimental bias. The low-noise design also allows for the integration of the NC4gate into neural recording applications. This open-source tool, and others like it, help to overcome resource limitations to enable greater innovation and integration of leading-edge approaches in rodent behavioral research.

